# Proton pump inhibitors inhibit PHOSPHO1 activity and matrix mineralisation *in vitro*

**DOI:** 10.1101/2021.04.29.441931

**Authors:** Katherine A. Staines, Katherine Myers, Kirsty Little, Stuart H. Ralston, Colin Farquharson

**Affiliations:** School of Pharmacy and Biomolecular Sciences, University of Brighton, Brighton UK; The Roslin Institute, The University of Edinburgh, Edinburgh, UK; Centre for Genomic and Experimental Medicine, Institute of Genetics and Cancer, University of Edinburgh, Edinburgh, United Kingdom

**Author notes:** Corresponding author*, Katherine Ann Staines, University of Brighton, Lewes Road, Brighton BN2 4GJ.

**Keywords:** PHOSPHO1, proton pump inhibitors, histamine-2 receptor antagonists, mineralisation, TNAP

## Abstract

Proton pump inhibitors (PPIs) have been associated with an increased risk of fragility fractures in pharmaco-epidemiological studies. The mechanism is unclear but it has been speculated that by neutralising gastric acid, they may reduce intestinal calcium absorption, causing secondary hyperparathyroidism and bone loss. Here we investigated that hypothesis that the skeletal effects of PPI might be mediated by inhibitory effects on the bone-specific phosphatase PHOSPHO1. We found that the all PPI tested potential inhibited the activity of PHOSPHO1 with IC50 ranging between 0.73μM for esomeprazole to 19.27μM for pantoprazole. In contrast, these PPIs did not inhibit TNAP activity. We also found that mineralisation of bone matrix in primary osteoblast cultures inhibited by several PPI in a concentration dependent manner. In contrast, the histamine-2 receptor antagonists (H2RA) nizatidine, famotidine, cimetidine and ranitidine had no inhibitory effects on PHOSPHO1 activity. Our experiments shown for the first time that PPI inhibit PHOSPHO1 activity and matrix mineralisation *in vitro* revealing a potential mechanism by which these widely used drugs are associated with the risk of fractures.

## Introduction

Proton pump inhibitors (PPIs) are amongst the most commonly prescribed drugs and are used in the treatment of gastroesophageal reflux disease (GORD), peptic ulcer disease and dyspepsia [1]. In the UK alone, more than 60 million PPI prescriptions were issued during 2017 [2]. The safety records of PPI’s are generally favourable but pharmaco-epidemiological evidence has consistently shown a positive association between PPI use and bone fractures. For example, large scale studies conducted in Denmark, UK and Canada all reported an increased risk of osteoporosis related fractures including fractures to the hip and spine with chronic PPI therapy [3–5].

The most commonly accepted explanation is that PPIs predispose to fractures by neutralising gastric acid. This in turn is thought to impair intestinal calcium absorption, secondary hyperparathyroidism and increased osteoclastic bone resorption with bone loss [6–8]. However, in healthy subjects, short term treatment with the PPI omeprazole was not found to have inhibitory effects on calcium absorption [9, 10]. Furthermore, epidemiological studies with histamine 2 receptor antagonists (H2RAs), which also supress gastric acid secretion, have not shown an association with fractures [3, 11–15]. Likewise, a recent meta-analysis reported that the use of PPIs, but not H2RAs, is associated with an increased risk of hip fracture [16]. These conflicting data suggest that PPI use may increase fracture incidence by a mechanism that distinct from effects on intestinal calcium absorption.

PHOSPHO1, a member of the haloacid dehalogenase superfamily, is a cytosolic phosphatase highly expressed by osteoblasts which is essential for bone mineralisation [17]. It liberates inorganic phosphate (P_i_) through the hydrolysis of phospholipid substrates within the matrix vesicle (MV) membrane [17–19]. Within this protected environment, Pi accumulates and chelates with Ca^2+^ which is enriched in MVs to form mineral crystals which subsequently invade and mineralise the organic collagenous scaffold [17–22]. Deletion of PHOSPHO1 in mice results in bowed long bones and spontaneous greenstick fractures, decreased cortical BMD and accumulation of osteoid in trabecular bone [23]. Similarly, osteoblasts treated with a PHOSPHO1 specific inhibitor and cultures of *Phospho1* deficient primary osteoblast both revealed reduced matrix mineralising ability, whereas matrix mineralisation was increased by osteoblasts overexpressing PHOSPHO1 [24, 25]. A critical role for PHOSPHO1 in the mineralisation process was confirmed in a comparison of the bone phenotype of; *Alpl*^-/-^; *Phospho1*^-/-^ double knockout mice to that of *Alpl*^-/-^ and *Phospho1*^-/-^ mice. The skeleton of both single gene knockouts was impaired whereas the double ablation led to the complete absence of skeletal mineralisation and embryonic lethality. These experimental data are consistent with the notion that PHOSPHO1 and TNAP have independent, non-redundant roles during the mineralisation process [23].

We previously identified, through a screen of chemical libraries containing over 50,000 compounds, the PPI, lansoprazole as a PHOSPHO1-specific inhibitor [18]. Indeed, lansoprazole non-competitively inhibited recombinant human PHOSPHO1 activity by over 70% and caused a 57% inhibition of osteoblast MV calcification but had no effect on tissue non-specific alkaline phosphatase (TNAP) activity [18]. Furthermore, *in vivo* studies disclosed that lansoprazole administration to developing chick embryos completely inhibited mineralisation of all leg and wing long bones [26].

In view of the fact that PHOSPHO1 plays a critical role in bone mineralisation, we hypothesise that the association between PPI use and bone fractures is possibly due to their inhibitory effect on PHOSPHO1 activity. To address this hypothesis, we used *in vitro* approaches to evaluate the potential of commonly prescribed PPIs and H2RAs to inhibit both PHOSPHO1 enzyme activity and osteoblast matrix mineralisation.

## Materials and Methods

### PPI and H2RAs

The PPIs lansoprazole, omeprazole, pantoprazole and esomeprazole (Cayman Chemicals, Michigan, USA) were used at varying concentrations (0-100μM) in the phosphatase activity and *in vitro* mineralisation assays detailed below. Similarly, the H2RAs nizatidine, famotidine, cimetidine and ranitidine (Selleckchem, Munich, Germany) were also used at 0- 100μM.

### Primary osteoblast isolation

Primary calvarial osteoblasts were obtained from 4-day-old wild-typeC57Bl/6 mice. All experimental protocols were approved by Roslin Institute’s Animal Users Committee and the animals were maintained in accordance with UK Home Office guidelines for the care and use of laboratory animals. Primary osteoblasts were isolated by sequential enzyme digestion of excised calvarial bones using a four-step process as has previously been described [7,8] [1 mg/ml collagenase type II in Hanks’ balanced salt solution (HBSS) for 10 min; 1 mg/ml collagenase type II in HBSS for 30 min; 4 mM EDTA for 10 min; 1 mg/ml collagenase type II in HBSS for 30 min]. The first digest was discarded and the cells were re-suspended in growth medium consisting of a-MEM (Invitrogen, Paisley, UK) supplemented with 10% (v/v) FBS and 1% gentamycin (Invitrogen). Osteoblasts were seeded at a density of 1 x 10^4^ cells/cm^2^ and grown to confluency at which point 2mM β-glycerophosphate and 50μg/ml ascorbic acid was added along with a PPI (0- 50μM) as described in results. Media was changed every 2-3 days for the duration of the 28-day experiments.

### Assessment of primary osteoblast matrix mineralisation

After 28 days, primary cell cultures were fixed in 4% paraformaldehyde for 5 min at room temperature. Cell monolayers were stained with aqueous 2% (w/v) Alizarin red solution for 5 min at room temperature. The bound stain was solubilised in 10% cetylpyridinium chloride and the optical density of the resultant eluted solution measured by spectrophotometry at 570nm.

### Phosphatase assays

Recombinant human PHOSPHO1 (50ng) was generated as previously described [27] and incubated with varying concentrations of the aforementioned PPIs and H2RAs in experimental assay buffer (20mM Tris, 2mM MgCl_2_ & 25μg/ml BSA) at 37°C for 15 mins. Using the BIOMOL^®^ Green assay (Enzo, Exeter, UK), standards (0-2nM) and samples were then incubated with 2.5mM β-glycerol phosphate for 30min at 37°C with gentle agitation [27]. The reaction was stopped using 100μl BIOMOL Green and after being left for 30min at room temperature, the absorbance was read using spectrophotometry at 630nm. For TNAP, 2ng recombinant human TNAP (R&D Systems, Abington, UK), was incubated with varying concentrations of the aforementioned PPIs and H2RAs in experimental assay buffer (1M diethylamine hydrochloride, 1mM MgCl_2_ and 20μM ZnCl_2_). Using the BIOMOL^®^ Green assay, standards (0-2nM) and samples were then incubated with 0.5mM p-nitrophenyl phosphate (pNPP) for 30min at 37°C with gentle agitation. The reaction was stopped using 100μl BIOMOL^®^ Green and after being left for 30min at room temperature, the absorbance was read using spectrophotometry at 630nm.

### Statistical analysis

Data are expressed as the mean ± standard error of the mean (S.E.M) of at least 3 replicates per experiment. Statistical analysis was performed by one-way analysis of variance (ANOVA). P<0.05 was considered to be significant and noted as *; P values of <0.01 and <0.001 were noted as ‘**’ and ‘***’ respectively.

## Results

### PPIs are potent inhibitors of PHOSPHO1 activity

In accordance with our previous results, lansoprazole inhibited PHOSPHO1 activity (IC_50_ = 2.767μM; Fig. 1A). Similarly, here we show for the first time that the PPIs omeprazole (IC_50_ = 2.803μM) and esomeprazole (IC_50_ = 0.726μM) are potent inhibitors of PHOSPHO1 activity (Figs. 1B & C). Whilst pantoprazole also inhibited PHOSPHO1 activity, its IC50 was 19.27μM, suggesting that this PPI is the least potent PHOSPHO1 inhibitor tested (Fig. 1D).

**Figure 1.**
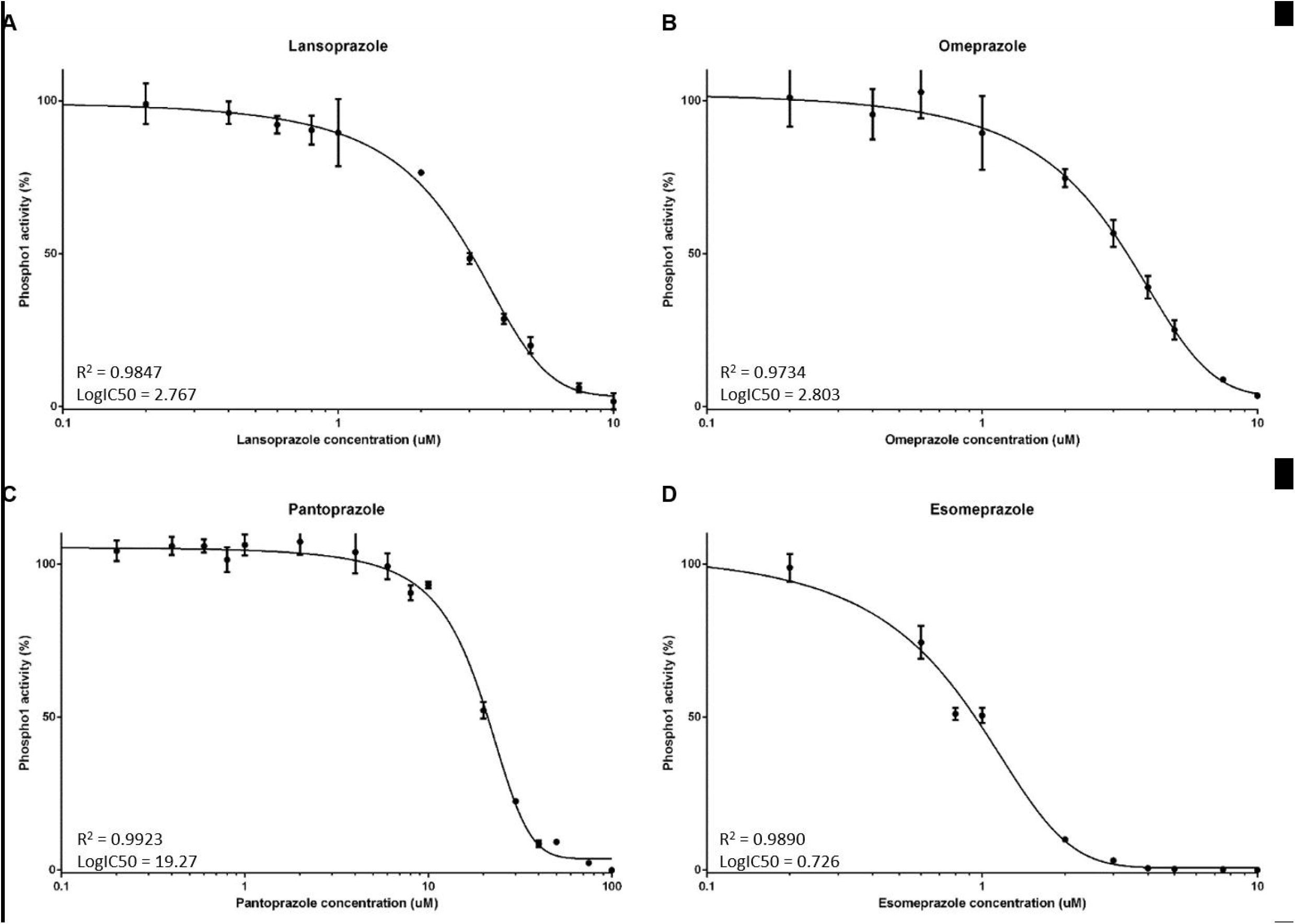
The effects of proton pump inhibitors (PPIs) on PHOSPHO1 activity. PHOSPHO1 activity was assessed by phosphatase assays in the presence of **(A)** lansoprazole **(B)** omeprazole **(C)** esomeprazole **(D)** pantoprazole.

### PHOSPHO1 activity is not inhibited by H2RAs

We next sought to examine whether PHOSPHO1 activity is similarly inhibited by four commonly prescribed H2RAs. At all concentrations tested, there was no inhibition of PHOSPHO1 activity upon addition of nizatidine (Fig. 2A), famotidine (Fig. 2B), cimetidine (Fig. 2C) and ranitidine (Fig. 2D).

**Figure 2.**
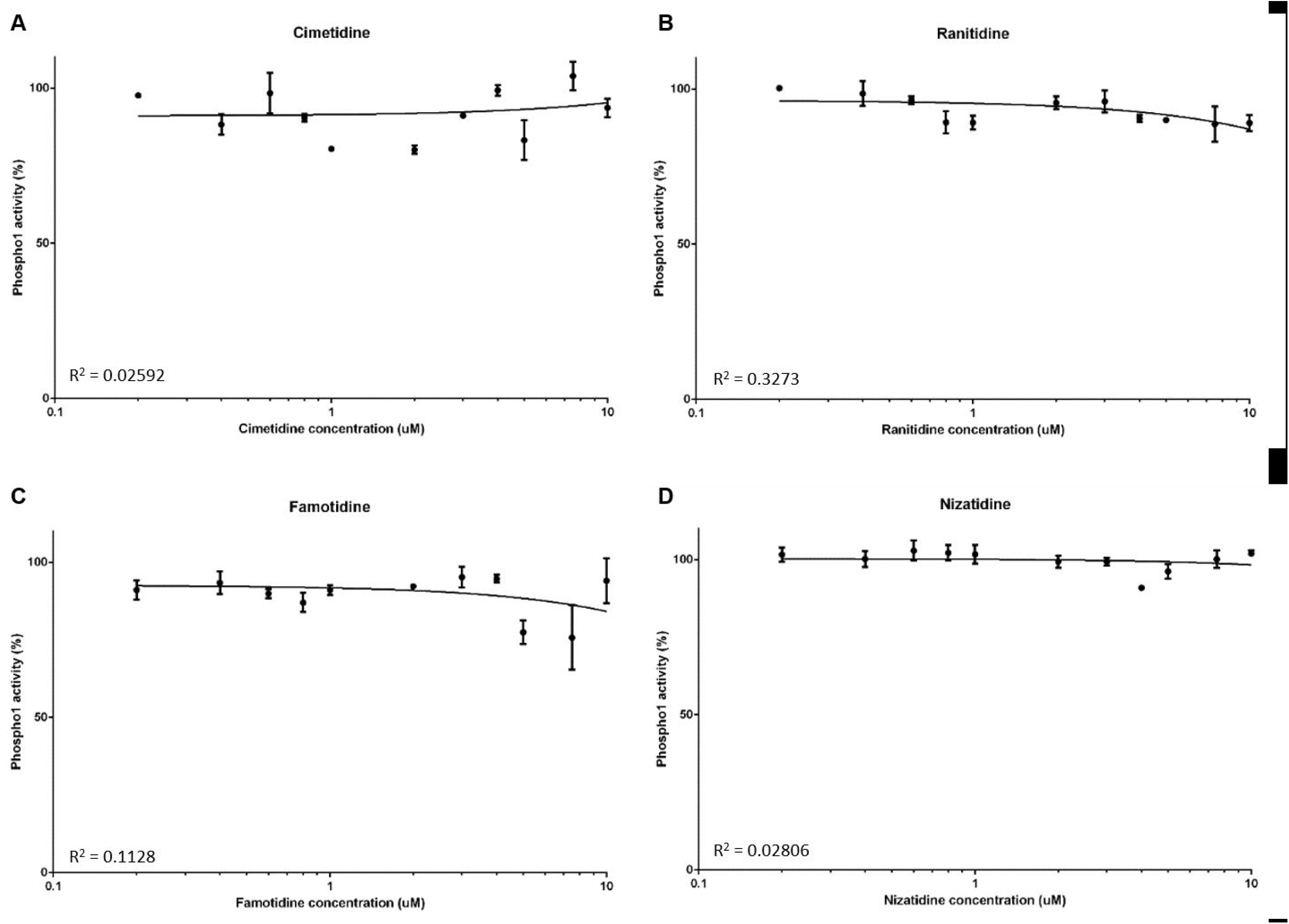
The effects of histamine-2 receptor antagonists (H2RAs) on PHOSPHO1 activity. PHOSPHO1 activity was assessed by phosphatase assays in the presence of **(A)** cimetidine **(B)** ranitidine **(C)** famitidine **(D)** nizatidine.

### PPIs and H2RAs have no effect on TNAP activity

We next determined whether the aforementioned PPIs are able to inhibit TNAP activity. At all concentrations tested, lansoprazole, omeprazole, esomeprazole and pantoprazole did not inhibit TNAP activity (Figs. 3A – D). Similarly, there was no inhibition of TNAP activity by the H2RAs (Fig. 4A – D).

**Figure 3.**
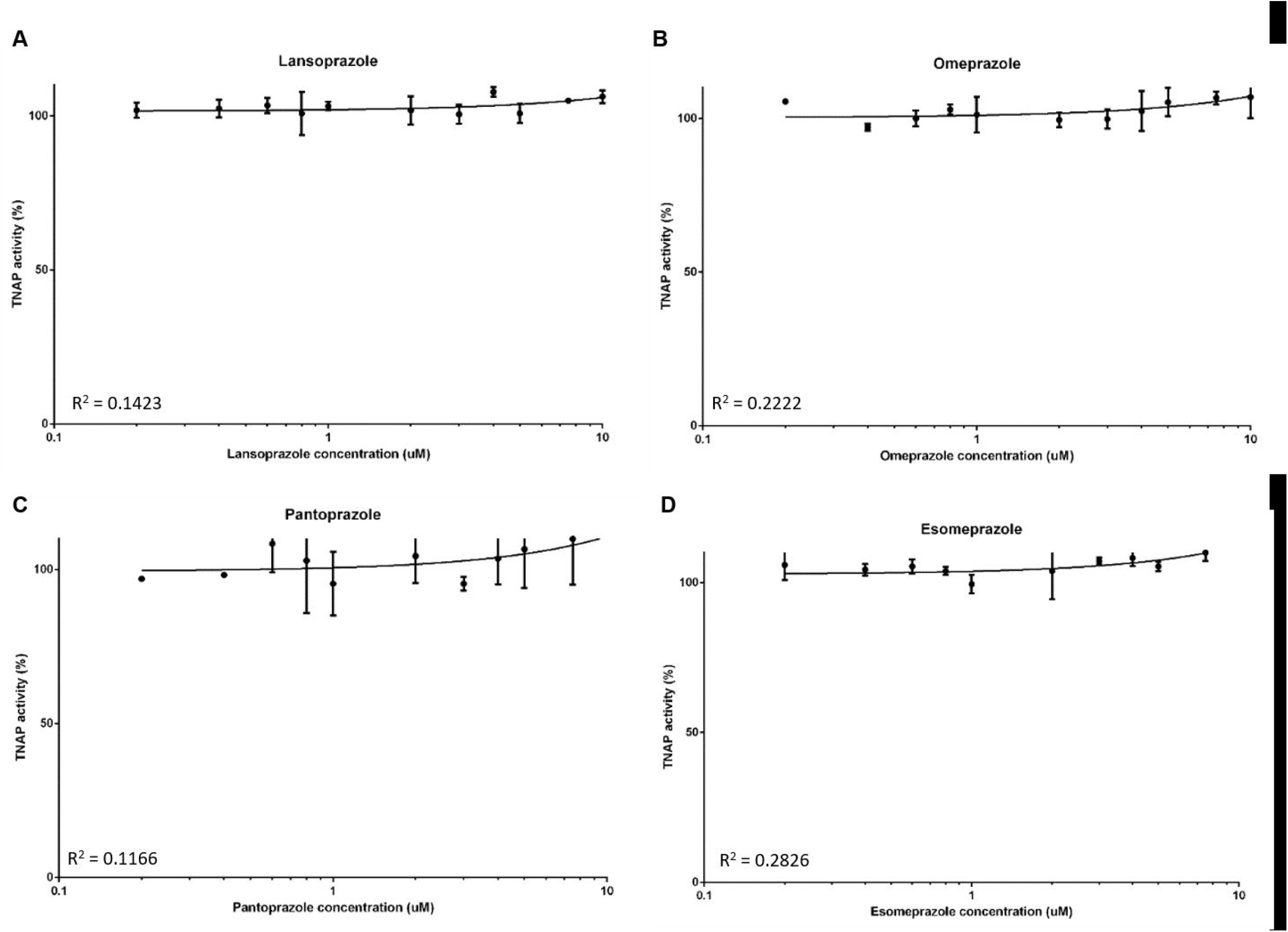
The effects of proton pump inhibitors (PPIs) on TNAP activity. TNAP activity was assessed by phosphatase assays in the presence of the PPIs **(A)** lansoprazole **(B)** omeprazole **(C)** esomeprazole **(D)** pantoprazole

**Figure 4.**
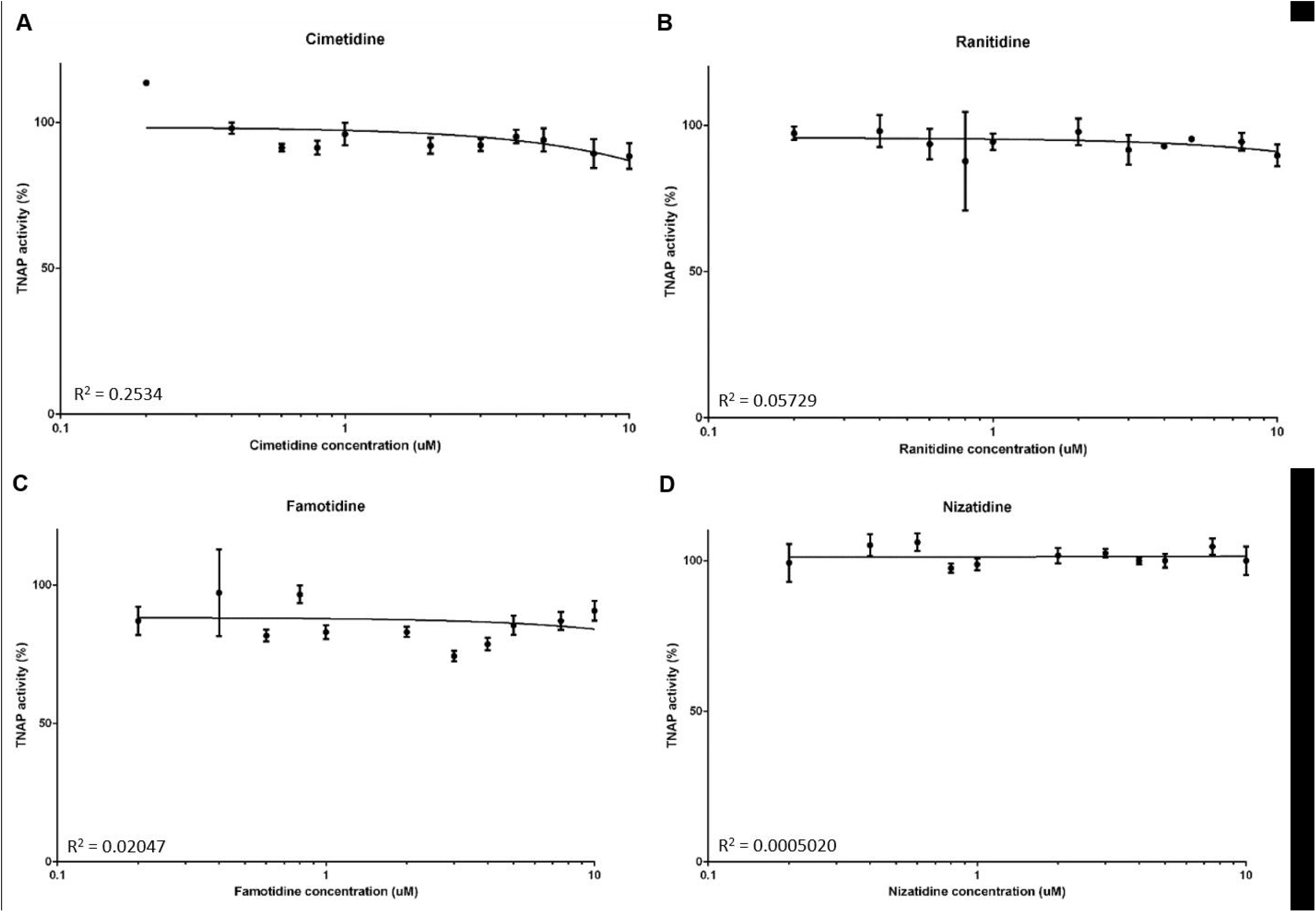
The effects of histamine-2 receptor antagonists (H2RAs) on TNAP activity. TNAP activity was assessed by phosphatase assays in the presence of the H2RAs **(A)** cimetidine **(B)** ranitidine **(C)** famitidine **(D)** nizatidine.

### PP1s inhibit primary osteoblast matrix mineralisation

To examine whether the inhibition of PHOSPHO1 by PPIs has an effect on matrix mineralisation, we cultured primary osteoblasts in the presence of different concentrations of lansoprazole, omeprazole, esomeprazole and pantoprazole. We found that whilst control cultures formed mineralised nodules after 28 days in culture, the addition of 5μM and 10μM lansoprazole significantly decreased matrix mineralisation (Figs. 5A, B & C). Despite this, nodules were clearly visible throughout the cultures suggestive that the effects seen are directly on mineralisation rather than the differentiation of the cells (Fig. 5A). Similarly, omeprazole and esomeprazole significantly inhibited matrix mineralisation at concentration of 10μM (Figs. 5A, B & C). In concordance with the higher IC_50_ of pantoprazole, culture of primary osteoblasts with 5μM and 10μM pantoprazole was not sufficient to inhibit matrix mineralisation (Figs. 5A, B & C). We therefore cultured cells with 50μM pantoprazole and indeed saw a significant decrease in matrix mineralisation (Figs. 5D).

**Figure 5.**
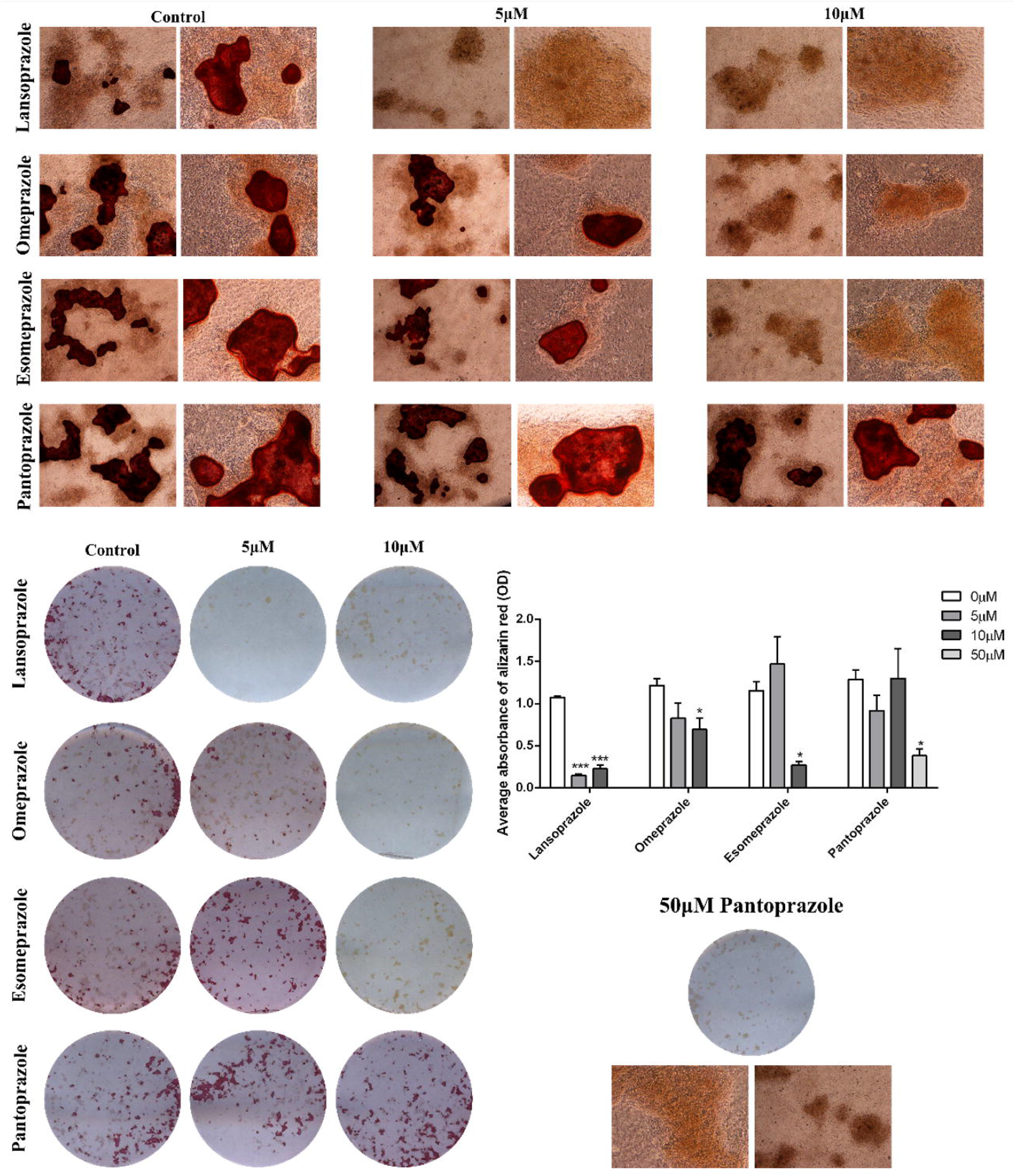
The effects of proton pump inhibitors (PPIs) on primary osteoblast matrix mineralisation. Primary osteoblasts were cultured for 28 days in the presence of 0-10μM lansoprazole, omeprazole, esomeprazole and pantoprazole. **(A)** Microscopic images of alizarin red stained mineral associated with nodule formation **(B)** Alizarin red staining **(C)** Quantification of alizarin red staining **(D)** Alizarin red staining of primary osteoblasts treated with 50μM pantoprazole. Data are represented as mean ± S.E.M. (n=3 wells/treatment) P<0.05*, P<0.01**, P<0.001***.

## Discussion

In this study we report that all the PPIs tested were inhibitors of PHOSPHO1 activity whilst they had no effect on TNAP activity. The most potent inhibitor was esomeprazole which gave 50% inhibition in the sub micromolar range, followed by lansoprazole, omeprazole and pantoprazole Consistent with this, the PPIs we tested inhibited mineralisation of bone matrix in vitro in low micromolar concentrations, except pantoprazole which did not have inhibitory effects until higher concentrations of 50uM were used. Conversely, we tested several H2RAs and these had no effect on PHOSPHO1 or TNAP phosphatase activity or on matrix mineralisation in vitro.

Several studies have shown an association with between PPIs use and fractures. Indeed, a large scale meta-analysis has reported a significant increase in relative risk (RR) of fractures at the hip [RR=1.26, 95% CI = 1.16-1.36] spine [RR=1.58, 95% CI = 1.38-1.82] and any-site fractures [RR=1.33, 95% CI = 1.15-1.54] in PPI users as compared with controls [29].

The PPIs reduce gastric acid secretion through inhibition of H+/K+-ATPases located in stomach parietal cells [28]. In view of this it has been speculated that calcium malabsorption mediated by neutralisation of gastric acid may cause secondary hyperparathyroidism and bone loss [6–8]. Other potential mechanisms include (i) impaired bone resorption resulting in altered bone remodelling and (ii) hypergastrinemia resulting in parathyroid hyperplasia and decreased bone mineral density [30, 31]. The H2RAs are also widely used to suppress gastric acid production in the treatment of GORD, dyspepsia and peptic ulcers these have not been associated with fractures in epidemiological studies which calls into question the hypothesis that the association between fractures and PPI used is mediated by reduced calcium absorption due to achlorhydria [3, 11–15, 33]. The = data presented here is consistent with this and suggests that inhibition of PHOSPO1 may be an alternative mechanism by which PPIs, affect bone health. The PHOSPHO1 enzyme is a bone specific phosphatase that is highly expressed at sites of mineralization and essential for the formation of mechanically competent bone [17]. It is biochemically active within MVs [18] and it has been proposed that the accumulation of Pi within MVs is a consequence of PHOSPHO1s intravesicular activity and also intravesicular trafficking of TNAP□generated Pi via a Type III Na□Pi co transporter, PiT1 [34–36]. We have previously shown that MV mineralisation is reduced in *Phospho1*^-/-^ mice [35, 37] and that lansoprazole treatment of MVs isolated from osteoblasts impairs their mineralisation [26]. It is therefore possible that PPI inhibition of PHOSPHO1 activity disrupts the biochemical machinery needed to establish the appropriate inorganic pyrophosphate to Pi ratio required to initiate the formation of HA mineral within MVs [36, 38]. Our *in vitro* cell culture work is also consistent with a previous study in which lansoprazole, esomeprazole and omeprazole decreased the ability of osteoblasts to mineralise their matrix, whilst also inhibiting osteoblast gene expression [39] These observations at the cell and MV level are consistent with, and explain, the reduced bone mineral content and BMD in rodents administered omeprazole [40, 41].

Interestingly, the data of this present study indicated no effect of PPIs on TNAP phosphatase activity; a result that is consistent with our previous study that reported lansoprazole and other small molecule inhibitors of PHOSPHO1 had no effect on TNAP activity [18]. The importance of TNAP in the mineralisation process is well accepted [42, 43]. Indeed, in patients with hypophosphatasia and also in *Alpl*^-/-^ mice, extravesicular crystal propagation is retarded due to an accumulation of inorganic pyrophosphate in the extracellular matrix [46]. These data imply that the inhibition of osteoblast matrix mineralisation by the PPIs is via their inhibition of PHOSPHO1, and not TNAP activity. A note of caution in the interpretation of these data is nevertheless warranted; other *in vitro* studies have reported that lansoprazole can inhibit porcine TNAP activity albeit with a Ki value of ~100 times higher than that reported for the inhibition of recombinant human PHOSPHO1 with lansoprazole [18, 50]. An explanation for these different results is unclear.

The order of potency (based on our IC50 data) of PPI inhibition of PHOSPHO1 activity is esomeprazole > omeprazole = lansoprazole > pantoprazole (Fig. 2), which precisely mimics our data in mineralising primary osteoblasts, but also their ability (based on omeprazole equivalents) to inhibit acid production [51, 52]. Intriguingly, this suggests that the structure of the more potent acid suppressive PPIs accounts for their PHOSPHO1 inhibitory properties. Also, pantoprazole, the PPI least able to inhibit PHOSPHO1 enzyme activity was also a poor inhibitor of matrix mineralisation. Knowing the molecular model of PHOSPHO1 [21], it would be of interest to perform ligand docking studies to gain more information as to how the different PPIs associate with the enzyme and temper its biological activity. This has the potential to equip industry with the knowledge to generate modified and improved PPIs without the undesired off target bone effects.

In summary we have shown that commonly prescribed PPIs, but not H2RAs, inhibit the activity of the bone specific phosphatase, PHOSPHO1 *in vitro* in a dose-dependent manner and at concentrations that are similar to those used clinically. We have also shown that different PPIs differ by more than 25-fold in their ability to inhibit PHOSPHO1 activity compared with a 7-fold difference in potency for inhibition of acid production [51]. This indicates that there is a >3-fold difference in the ability of PPIs to inhibit PHOSPHO1 activity as compared with their ability to suppress gastric acid production.

In view of the fact that PHOSPHO1 plays a critical role in bone mineralisation, we hypothesise that the association between PPI use and bone fractures is possibly due to their inhibitory effect on PHOSPHO1 activity. While this remains to be confirmed by further research it could have clinical implications in allowing clinicians to select PPI’s with the least inhibitory effect on PHOSPHO1 activity as the preferred drug in this class in patients at high risk of fragility fractures.

## Availability of data

Data are available on reasonable request from the corresponding authors.

## Authors contributions

Conception and design of the study: KS, SR, CF. Acquisition of data: KS, KM, KL. Interpretation of data: KS, CF. Drafting the manuscript: KS, CF. Revising the manuscript and final approval, and agreement to be accountable for all aspects of the work: all authors.

## Ethics approval

All experimental protocols were approved by Roslin Institute’s Animal Users Committee and the animals were maintained in accordance with UK Home Office guidelines for the care and use of laboratory animals.

